# Genetic Architecture of Kernel Compositional Variation in a Maize Diversity Panel

**DOI:** 10.1101/2021.03.29.436703

**Authors:** Jonathan S. Renk, Amanda M. Gilbert, Travis J. Hattery, Christine H. O’Connor, Patrick J. Monnahan, Nickolas Anderson, Amanda J. Waters, David P. Eickholt, Sherry A. Flint-Garcia, Marna D. Yandeau-Nelson, Candice N. Hirsch

**Affiliations:** Department of Agronomy and Plant Genetics, University of Minnesota, St. Paul, MN 55108; Department of Genetics, Development, and Cell Biology, Iowa State University, Ames, IA 50011; Department of Ecology, Evolution, and Behavior, University of Minnesota, St. Paul, MN 55108; PepsiCo R&D, St Paul, MN 55108; United States Department of Agriculture, Agricultural Research Service, Columbia, MO 65211

## Abstract

Maize (*Zea mays* L.) is a multi-purpose row crop grown worldwide, which overtime has often been bred for increased yield at the detriment of lower composition grain quality. Some knowledge of the genetic factors that affect quality traits has been discovered through the study of classical maize mutants. However, much of the underlying genetic architecture controlling these traits and the interaction between these traits remains unknown. To better understand variation that exists for grain compositional traits in maize, we evaluated 501 diverse temperate maize inbred lines in five unique environments and predicted 16 compositional traits (e.g. carbohydrates, protein, starch) based on the output of near-infrared (NIR) spectroscopy. Phenotypic analysis found substantial variation for compositional traits and the majority of variation was explained by genetic and environmental factors. Correlations and trade-offs among traits in different maize types (e.g. dent, sweetcorn, popcorn) were explored and significant differences and correlations were detected. In total, 22.9-71.1% of the phenotypic variation across these traits could be explained using 2,386,666 single nucleotide polymorphism (SNP) markers generated from whole genome resequencing data. A genome-wide association study (GWAS) was conducted using these same markers and found 70 statistically significant loci for 12 compositional traits. This study provides valuable insights in the phenotypic variation and genetic architecture underlying compositional traits that can be used in breeding programs for improving maize grain quality.

**Core Ideas:**

1. Understanding kernel compositional variation is important for food grade corn improvement.
2. Genetic and environmental factors account for most of the variation in compositional traits.
3. A broad range in trait heritabilities was observed across compositional traits.
4. Compositional trade-offs will be important to consider when conducting multitrait breeding.
5. Compositional traits are mostly controlled by a large number of small effect loci.

## INTRODUCTION

Maize (*Zea mays* L.) is a multi-purpose crop with a wide variety of uses including feed, fuel, food, and industrial products. In the United States, the vast majority of grain harvested does not contribute directly to human consumption or food products. In 2019, it was estimated that only around 3.5% of the grain harvested was used for cereals and other food products (USDA NASS, 2019). In the United States, demand and consumption of maize-based food products has tripled since the 1970s, primarily through tortillas and tortilla chips due to the popularity of both Hispanic and gluten-free food products (Scott et al., 2019). Maize is also a staple food crop for many other regions around the globe including Latin America and Africa. To meet increasing market demands in the United States and globally, the focus on grain quality in maize breeding programs has become paramount.

Compositional traits are important factors in determining the overall quality of the grain and in turn food production. However, in the United States much of the focus of maize breeding has been on yield gain, while neglecting quality traits through much of the breeding pipeline (Holmes et al., 2019). This is quite evident when looking at kernel composition over time in the United States. Investigating commercial hybrids from 1960 to 1990, Duvick and Cassman (1999) found that grain protein content has steadily decreased and starch content has increased in a trade-off to select higher yielding hybrids. This change is a negative consequence for the cooking process known as nixtamalization, in which maize kernels are cooked in an alkaline solution to facilitate the removal of the pericarp and are ground into masa dough (Serna-Saldivar & Rooney, 2015). For example, dry matter loss (DML) during cooking is a very important trait to commercial producers as it adds to the production cost of making food products. Pflugfelder et al., (1988) found that hybrids with softer crown and endosperm in the kernel due to higher starch content have a higher amount of DML. The reduction in kernel hardness can also cause a higher level of kernel cracking or stress cracks during storage and drying of the grain at harvest, which leads to increased moisture uptake in the kernel (Jackson et al., 1988), with subsequent impacts on final product quality.

To shift the focus of breeding programs to improved quality traits as a primary breeding objective, a deep understanding of the genetic factors that contribute to variation for compositional traits is beneficial. Much of the current knowledge on the genetics of quality traits comes from the study of classical mutants in maize. Some examples of known mutants include *sugary1* (*su1*) (James et al., 1995), *sugary2* (*su2*) (Zhang et al., 2004), and *waxy1* (*wx1*) (Tsai, 1974), which are involved in the starch biosynthesis pathway and are critical for sweet corn breeding. The *opaque2* (*o2*) (Mertz et al., 1964), *floury2* (*fl2*) (Nelson et al., 1965), and *floury1* (Holding et al., 2007) mutants affect the amino acid content and zein protein composition in the kernel. The *o2* mutant has been used in quality protein maize (QPM) breeding programs to increase lysine and tryptophan content in animal feed from maize, with much of the work being done by the International Maize and Wheat Improvement Center (CIMMYT) (Babu et al., 2005). The utilization of these mutations has been difficult to deploy due to the negative effects on yield (Prasanna et al., 2001). Furthermore, long-term studies such as the Illinois Long-Term Selection experiment on oil and protein content in kernels suggests the genetic architecture of these traits is quite complex with many small effect loci contributing to phenotypic variation (Dudley & Lambert, 2004).

Genome-wide association study (GWAS) is a powerful tool that allows researchers to identify regions of the genome that correlate with a trait of interest and determine the genetic architecture of a trait within a particular germplasm set. The resolution of association mapping depends on the historical recombination that overtime produces smaller linkage blocks that can be associated with traits (Zhu et al., 2008). A limited number of GWAS have been performed for kernel compositional traits in maize. For example, protein, starch, and oil estimates generated from near-infrared (NIR) spectroscopy prediction equations in the maize nested association mapping (NAM) population (McMullen et al., 2009; Yu et al., 2008) and a panel of 302 diverse inbred lines (Flint-Garcia et al., 2005) found 26, 21, and 22 quantitative trait loci (QTL) respectively through joint-linkage mapping (Cook et al., 2012). A study of oil and fatty acid composition measured in the AM508 diversity panel of tropical, subtropical, and temperate inbred lines (Yang et al., 2011) revealed 74 QTL underlying these traits (Li et al., 2013). A subsequent study of this same diversity panel detected 247 and 281 statistically significant QTL for 17 proteinogenic amino acid traits in two separate environments (Deng et al., 2017). Finally, the genetic architecture of carotenoid-related traits was explored in CIMMYT’s 380 inbred carotenoid association panel and 40 QTL were identified for these traits (Suwarno et al., 2015) and Wang et al. (2020) identified *vitreous endosperm 1* (*ven1*) as a major QTL for influencing vitreous endosperm in the kernel.

Although there have been a handful of studies examining compositional traits in large germplasm panels, there is still a need for better understanding of the compositional trade-offs between traits, as well as their underlying genetic architecture in order to breed for enhanced composition important for human consumption and food products. Here, we evaluated the Wisconsin Diversity (WiDiv) population (Hansey et al., 2011; Hirsch et al., 2014; Mazaheri et al., 2019), across five environments and evaluated 16 compositional traits obtained from NIR spectroscopy prediction equations to investigate: 1) compositional trade-offs within the population, 2) the role of genotype by environment (GxE) interaction in trait variation, 3) how variation partitions differently across sets of germplasm that are adapted to grow in the Upper Midwest, and 4) the genetic architecture of these traits.

## MATERIALS AND METHODS

### Germplasm and Genomic Data

A subset of 501 lines from the full WiDiv population was used for this study (Supplemental Table 1). These lines all have available whole genome resequencing data with approximately 20x sequence depth, and SNPs from these data were previously identified (O’Connor et al., 2020). Briefly, reads were aligned to the B73 v4 reference genome assembly (Jiao et al., 2017) and joint SNP calling was performed using *freebayes* v1.3.1-17 (Garrison & Marth, 2012).

The SNP calls generated from *freebayes* were converted to HapMap format with the software package TASSEL version 5.2.64 (Bradbury et al., 2007). Within TASSEL, an overall genetic summary of genotype and site information was obtained. A pairwise genetic distance matrix was calculated by subtracting identity by state (IBS) from 1 to create a dendrogram that was exported in Newick Tree format, which was visualized using Figtree version 1.4.4 (Cummings, 2004). The resulting dendrogram was visually inspected for consistency with known pedigree relationships and shared heterotic groups. Samples with greater than 15% heterozygosity and loci with greater than 15% heterozygosity and 25% missing data were removed using the TASSEL filter menu for sites and taxa. This pruning resulted in the removal of six genotypes (PHJ90, Pa778, N534, 4N506, Va22, PHR63) and 733,462 SNPs, resulting in a 2,412,791 final SNP set (Supplemental File 1).

The SNP calls from the genomic resequencing data were compared to previous RNA-seq generated SNPs on the WiDiv association panel (Hirsch et al., 2014; Mazaheri et al., 2019). Pairwise genetic distance matrices were made for both datasets of common genotypes. A scatterplot comparing the genetic distances of the datasets was utilized to identify genotypes with inconsistencies between these two data sets (Supplemental Figure 1). Four additional samples (PHR55, B91, PHG86, B14A) were removed following this criteria. A dendrogram with this final set of genotypes and SNPs was once again generated using the same methods described above (Supplemental Figure 2).

### Field Experimental Design and Phenotypic Data Collection

The 501 genotypes were grown in field trials in the summers of 2016 and 2017. All experiments were planted as a randomized complete block design with two replications at each location. Within replicates, genotypes were further blocked by flowering time. The first block consisted of the earlier flowering lines (flowering at approximately 71 to 80 days after planting based on a previous planting of this population in St. Paul, MN), while the second block consisted of the later flowering lines (flowering at approximately 80 to 87 days after planting). Inbred lines B73 and PH207 were planted as checks within each block, with five entries of each check per block. Plots were grown as single rows (3.35m long and 0.76m apart) at approximately 70,000 plants ha^-1^. In the summer of 2016, trials were grown at the Minnesota Agricultural Experiment Station (St. Paul, MN) and the Iowa State University Agricultural Engineering/Agronomy Research Farm (Boone County, IA). The trial in St. Paul was planted on May 17, and ears were harvested between October 19-25. The trial in Boone County was planted on May 16, and ears were harvested between October 16-19. In 2017, trials were grown again at the Minnesota Agricultural Experiment Station, the Iowa State University Agricultural Engineering/Agronomy Research Farm, and also at the Bradford Research Center (Columbia, MO). The trial in St. Paul was planted on May 11, and ears were harvested between October 24-25. The trial in Boone County was planted on May 8, and ears were harvested on September 27 and October 4. The trial in Columbia was planted on May 15, and ears were harvested on September 28 and October 12.

Ears from ten representative plants in the middle of each plot were harvested and a random sample of 120 grams of kernels was ground using a Mill Feeder 3170 (Perten Instruments, Springfield, IL). Ground samples were scanned using a Perten DA 7250 near-infrared (NIR) spectroscopy scanner and global prediction equations, developed by Perten Instruments (www.perten.com), were used to predict Ash As Is, Fat As Is, Fiber As Is, Moisture, Protein As Is, and Starch As Is. As Is is defined as the trait value without correction for grain moisture at the time of the test. Local equations were also developed for the Perten DA 7250 NIR scanner for ten additional traits with wet chemistry run at Eurofins Scientific, Inc. (Des Moines, IA) using AOAC and AOCS standard protocols: Ash (AOAC 942.05), Ankom Crude Fiber (AOCS Ba 6a-05), Crude Fat (AOAC 920.39), Crude Fiber (AOAC 962.09; AOCS Ba 6-84), Fructose (AOAC 982.14), Glucose (AOAC 982.14), N Combustion (AOAC 992.15; AOAC 990.03; AOCS Ba 4e-93), Kjeltec Nitrogen (hereafter referred to as N Kjeltec; AOAC 2001.11), Sucrose (AOAC 982.14), and Total Sugars (AOAC 982.14) (AOAC, 2019; AOCS, 2020). Ankom crude fiber was measured by an Ankom^TM^ fiber analyzer. Each sample was run in triplicate for equation development. These equations were developed using a set of 100 spectrally diverse inbred lines from the WiDiv association panel (Supplemental Table 2). Wet lab data were calibrated to the Perten DA 7250 NIR scanner via the Honigs regression method within the Perten NIR software (Supplemental Figure 3) (Honigs et al., 1985). Raw data for the six global equations and ten local equations on the 4,886 plots are contained in Supplemental Table 3. It should be noted that for both the global and local equations some negative values are observed due to the fact that these values are predictions from regression equations.

### Statistical Analysis

Statistical analyses were conducted using R version 4.0.2 (R Core Development Team, 2020). Raw data were plotted for each trait as a whole and by environment to visually identify outliers. Extreme outliers were removed from the dataset if points were beyond five standard deviations from the mean of a given trait, which resulted in the removal of two data points from downstream analyses and from Supplemental Table 3. Each compositional trait was then analyzed using a random effects model *y = μ = g = e = r(e) = b(r) = gxe = ε* where *y* is the phenotypic response, *g* is the genotypic effect of an inbred line, *e* is the environmental effect of the year by location combination, *r(e)* is the effect of replicate within environment, *b(r)* is the effect of block within replicate, *gxe* is the genotype-by-environment interaction and *ε* is the residual. Models were fit with the restricted maximum likelihood method (REML) using the R package lme4 (Bates et al., 2014), and no violations were detected while checking model assumptions (independence of residuals, residual normality, and homoscedastic variance of residuals) (Supplemental Figure 4).

Broad sense heritability was calculated for each trait on an entry-mean basis using the following equation: 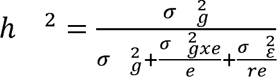 where 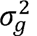 is the genotypic variance, 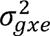 is the genotype-by-environment interaction variance, 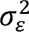 is the error variance, *e* is the number of environments, and *r* is the number of replications. Pearson correlations between NIR compositional predictions were calculated using the cor function in R. Correlations were considered statistically significant with a p-value <0.05. Correlations were conducted on the data set as a whole and then partitioned out based on heterotic groups and types of corn (e.g. dent, sweet corn, popcorn).

### Genome-wide Association Analysis

To investigate whether compositional traits should be analyzed separately for each environment or combined across environments, Spearman rank correlations were calculated between environments. Spearman rank correlations revealed that the environments were significantly correlated with each other (p-value < 0.05), and thus environments were combined. Best linear unbiased predictions (BLUPs) were extracted as the genotypic random effect from the model for each trait. All BLUPs can be found in Supplemental Table 4.

The Hapmap data generated in TASSEL was transformed into numerical format in R version 4.0.2 (R Core Development Team, 2020) using the package GAPIT version 3 (Lipka et al., 2012) by setting the numerical function flag equal to TRUE. GAPIT was also used to generate principal components to be used as covariates in GWAS (Supplemental Figure 5). The GWAS for the WiDiv BLUPs was conducted in R version 4.0.2 (R Core Development Team, 2020) utilizing the multiple locus linear mixed model (MLMM) in the package FarmCPU version 1.02 (Liu et al., 2016). In the model, the first five principal components were included as covariates to account for population structure (Supplemental Figure 5 and Supplemental Table 1), and a threshold for initial SNP inclusion into the model was set for each trait by allowing a false entry rate of 1% based on 100 permutations. SNPs with minor allele frequency less than 0.05 were discarded, resulting in a final SNP total for the GWAS analysis of 2,386,666. Optimal bin size and quantitative trait nucleotide (QTN) number were selected for each trait by selecting “optimum” under the “method.bin” parameter. SNPs were deemed statistically significant with a Bonferroni-adjusted p-value of <0.05. After excluding samples with high missingness, 446 samples (out of the initial 501) remained.

FarmCPU was chosen as the preferred method to conduct GWAS for a few reasons. First, a major issue with GWAS is that many false positives and false negatives appear in the results. To mitigate this issue, kinship and population structure of the association panel are often used as covariates in a Mixed Linear Model (MLM) to control false positives, which in turn weakens the signals of the associated SNPs. FarmCPU uses a multiple loci linear mixed model (MLMM) that incorporates multiple markers as covariates in a stepwise MLM to limit confounding markers and kinship (Liu et al., 2016). A complete elimination of any confounding from markers and kinship is achieved through the use of a MLMM in two parts in an iterative fashion with a fixed effects model (FEM) and a random effects (REM) model, which result in increased statistical power, computational efficiency, and control for both false positives and negatives (Liu et al., 2016). Kaler et al., (2020) conducted a study of simulated soybean and maize data comparing different statistical models used in GWAS and found that FarmCPU was able to control for false positives and false negatives and was consistently able to identify statistically significant SNPs closest to known published genes. It was also concluded that FarmCPU is a robust model for GWAS of complex traits in plants (Kaler et al., 2020).

Candidate genes for statistically significant SNP hits corresponded to physical positions based on annotation of the AGPv4 reference assembly of inbred B73 (Jiao et al., 2017). Functional annotations of candidate genes were obtained from MaizeGDB (Andorf et al., 2016), BLAST alignments to *Z. mays* (Madden, 2013), and Pfam domains (Finn et al., 2016).

Phenotypic variance explained (PVE) by the 2,386,666 SNPs used for GWAS was calculated as previously described using the software GCTA version 1.93 (Yang et al., 2010). Briefly, in TASSEL version 5.2.64 (Bradbury et al., 2007), SNPs were forced into a biallelic state and then the HapMap format was exported into PLINK ped and map files. In PLINK version 1.07 (Purcell et al., 2007), the two files were used to create the binary PED files used in GCTA. In GCTA, the GREML pipeline was used to calculate the genetic relationship matrix for all SNPs and PVE was estimated for all 16 traits based on the BLUPs generated from the random effects model as the phenotypic input file. In GREML, SNPs are fit jointly in a random-effect model and the only estimates of the model are the variance components, which is much smaller than the sample size so there is no overfitting.

### Code Availability

Code for statistical analysis and GWAS can be found at https://github.com/HirschLabUMN/GWAS_composition.

## RESULTS AND DISCUSSION

### Substantial variation for compositional traits exists within the WiDiv association panel

NIR predictions for 16 traits were collected for 501 inbred lines in the WiDiv association panel grown in two environments in 2016 (MN and IA) and three environments in 2017 (MN, IA, and MO). Visualization and descriptive statistics of raw values for compositional traits of the association panel show substantial variation for each trait roughly following normal distributions (Table 1 and Figure 1). The variation in compositional traits across this panel of lines is consistent with many other traits that have been previously measured in the WiDiv association panel such as flowering time (Hansey et al., 2011), tassel morphology (Gage et al., 2018), and stalk characteristics (Mazaheri et al., 2019).

**Figure 1.**
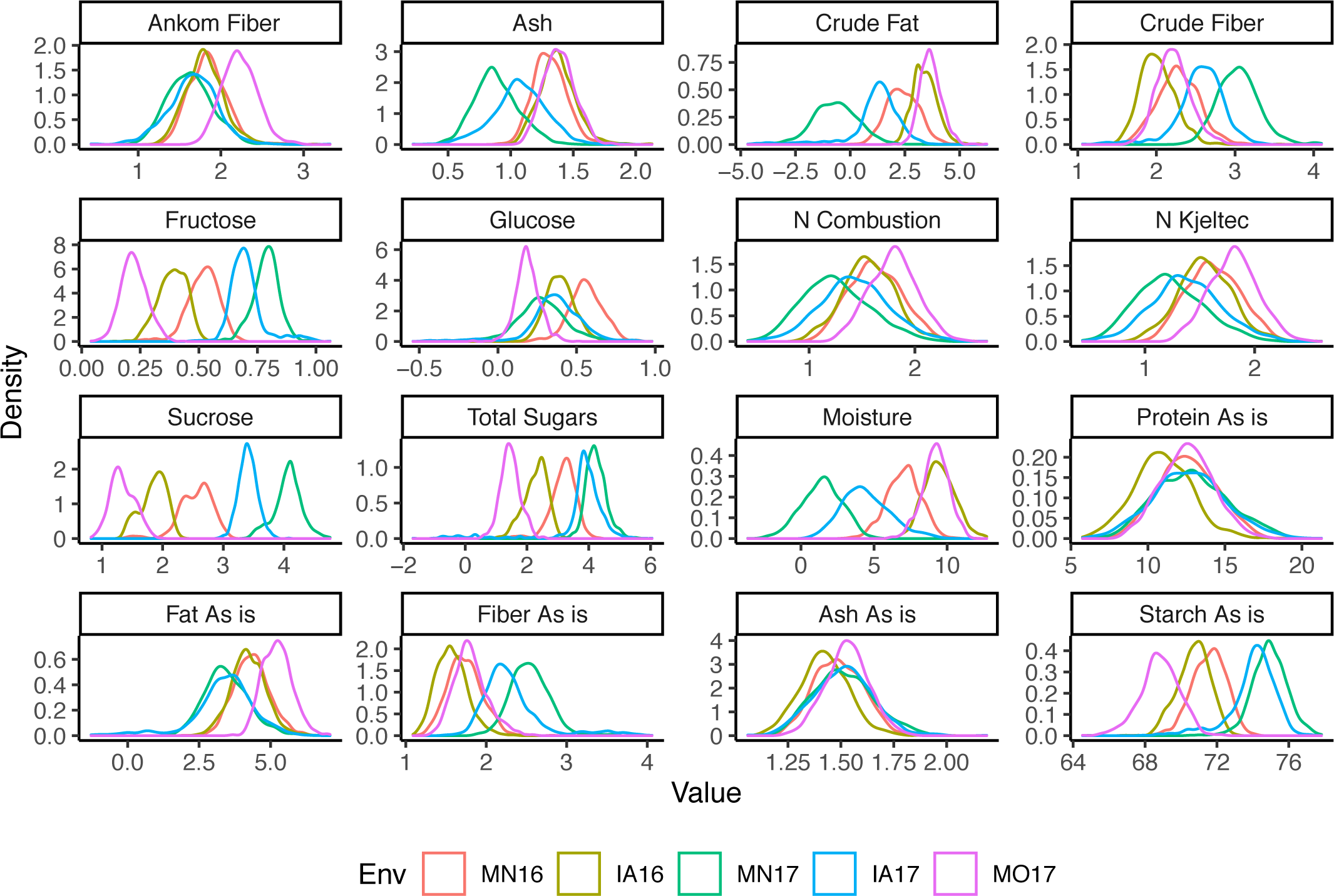
Trait distributions. All sixteen NIR prediction trait raw value distributions in each of the five unique environments in 2016 and 2017. The density plot lines are colored according to the corresponding environment. Env, environment; MN16, St. Paul, MN 2016; IA16, Ames, IA 2016; MN17, St. Paul, MN 2017; IA17, Ames, IA 2017; MO17, Columbia, MO 2017.

**Table 1:** Summary of traits and heritabilities.

| Trait | Mean (%) ± SD<sup>1</sup> | Range (%) | Heritability |
| --- | --- | --- | --- |
| Ankom Crude Fiber | 1.83±0.32 | 0.43-3.32 | 0.92 |
| Ash As Is | 1.49±0.13 | 1.06-2.19 | 0.94 |
| Ash | 1.21±0.25 | 0.22-2.13 | 0.89 |
| Crude Fat | 1.99±1.80 | (-)4.68-6.24 | 0.78 |
| Crude Fiber | 2.42±0.44 | 1.06-4.1 | 0.83 |
| Fat As Is | 4.17±1.00 | (-)1.28-7.08 | 0.88 |
| Fiber As Is | 1.99±0.45 | 1.09-4.06 | 0.84 |
| Fructose | 0.52±0.21 | 0.04-1.06 | 0.45 |
| Glucose | 0.36±0.18 | (-)0.54-0.98 | 0.54 |
| Moisture | 6.31±3.31 | (-)3.64-12.71 | 0.82 |
| Nitrogen Combustion | 1.53±0.33 | 0.42-2.68 | 0.94 |
| Nitrogen Kjeltec | 1.52±0.33 | 0.43-2.61 | 0.94 |
| Protein As Is | 12.32±2.17 | 5.75-21.27 | 0.94 |
| Starch As Is | 72.01±2.43 | 64.52-77.82 | 0.75 |
| Sucrose | 2.65±1.01 | 0.82-4.76 | 0.36 |
| Total Sugars | 3.00±1.13 | (-)1.69-6.05 | 0.49 |

<sup>1</sup>SD, standard deviation

Means, standard deviations, and ranges were calculated from the raw data across all environments (Table 1). Crude fat had the largest coefficient of variation (CV) (0.90) with values ranging from -4.68 to 6.24%. The smallest CV observed was for starch as is (0.03) with a range of 64.52 to 77.82%. For the global equation traits, protein as is ranged from 5.75 to 21.27%, fiber as is ranged from 1.09 to 4.06%, fat as is ranged from -1.28 to 7.08%, starch as is ranged from 64.52 to 77.82%, and ash as is ranged from 1.06 to 2.19% (Table 1). The ranges for the global equation traits in the WiDiv association panel are consistent with a previous study that analyzed a 282 inbred panel, in which starch, protein, and oil ranged from 59.6 to 70.3%, 11.5 to 17.5%, and 3.1 to 8.2%, respectively (Cook et al., 2012). Compared to a study by Reynolds et al., (2005) who analyzed seven contemporary maize hybrids the proximate starch content was higher than in WiDiv (79.3 to 88.0% compared to 64.52 to 77.82% in WiDiv), and also had a smaller range in protein (8.03 to 15.6% compared to 5.75 to 21.27% in WiDiv). This observation is consistent with a general trade-off between starch and protein content that has been observed in maize breeding history (Duvick & Cassman, 1999).

The 501 inbred lines of the WiDiv association panel consist of many different types of maize, including 145 Stiff Stalk, 143 Non-Stiff Stalk, 80 unknown, 54 mixed, 50 Iodent, 12 tropical, 11 popcorn, 5 sweet corn, and 1 flint based on pedigree information (Supplemental Table 1). Given the range of types of maize that comprised the subset of the WiDiv panel included in this study, we investigated the differences in composition between types and heterotic groups. BLUP-adjusted means were generated from a random effects model for each type and a Tukey’s HSD (*α* = 0.05) was used to investigate if statistically significant differences existed for compositional traits among maize types (Table 2). For starch as is, sweet corn had the lowest value at 70.87% and differed statistically from all other types. This can be attributed to the physiological differences among sweet corn kernels compared to dent kernels, in which sweet corn kernels have been bred to have increased sugar content and less starch content at maturity (Tracy, 2010). Popcorn had the highest amount of Ankom crude fiber, crude fiber, and fiber as is and differed statistically from the dent germplasm, i.e. Stiff Stalk, Non-Stiff Stalk and Iodent (Table 2). The majority of the fiber in corn kernels is found in the pericarp, which is the outermost layer that encases the entire kernel except the tip cap, and is composed largely of hemicelluloses including xylans and arabinoxylans (Chateigner-Boutin et al., 2016; García-Lara et al., 2019). In a study comparing anatomical percentages of kernel anatomy between dent, flint, and popcorn, popcorn had the highest proportion of pericarp at 7% followed by flint at 6.5% then dent at 6.0% (Serna-Saldivar, 1996). Stiff Stalk and Iodent types differed statistically for crude fat from tropical and for fat as is from tropical and sweet corn types, with the lowest values compared to the other maize types. In the kernel, the majority (∼85%) of fatty acids are found in the embryo (Shen & Roesler, 2017). This finding suggests that Stiff Stalk and Iodent types may have been selected and bred over time to have a smaller proportion of germ (i.e. embryo) and less protein in the endosperm, though not significantly different, which would increase starch content at the cost of protein within the endosperm compared to other maize types. No statistically significant differences between BLUP adjusted means for ash, nitrogen combustion, nitrogen Kjeltec, and protein as is were observed across maize types (Table 2).

**Table 2.**
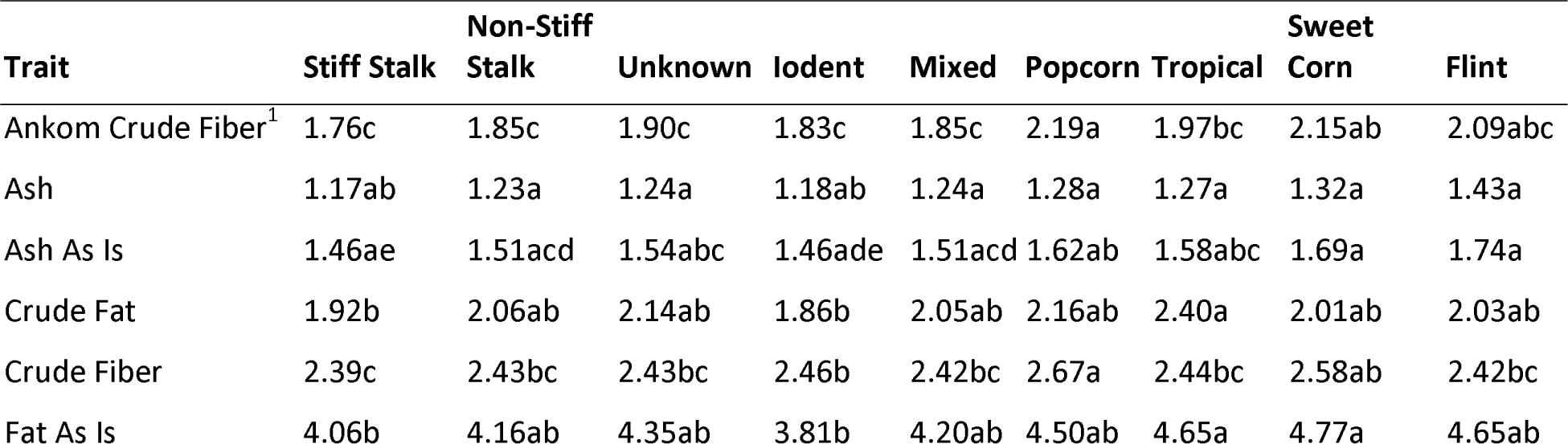
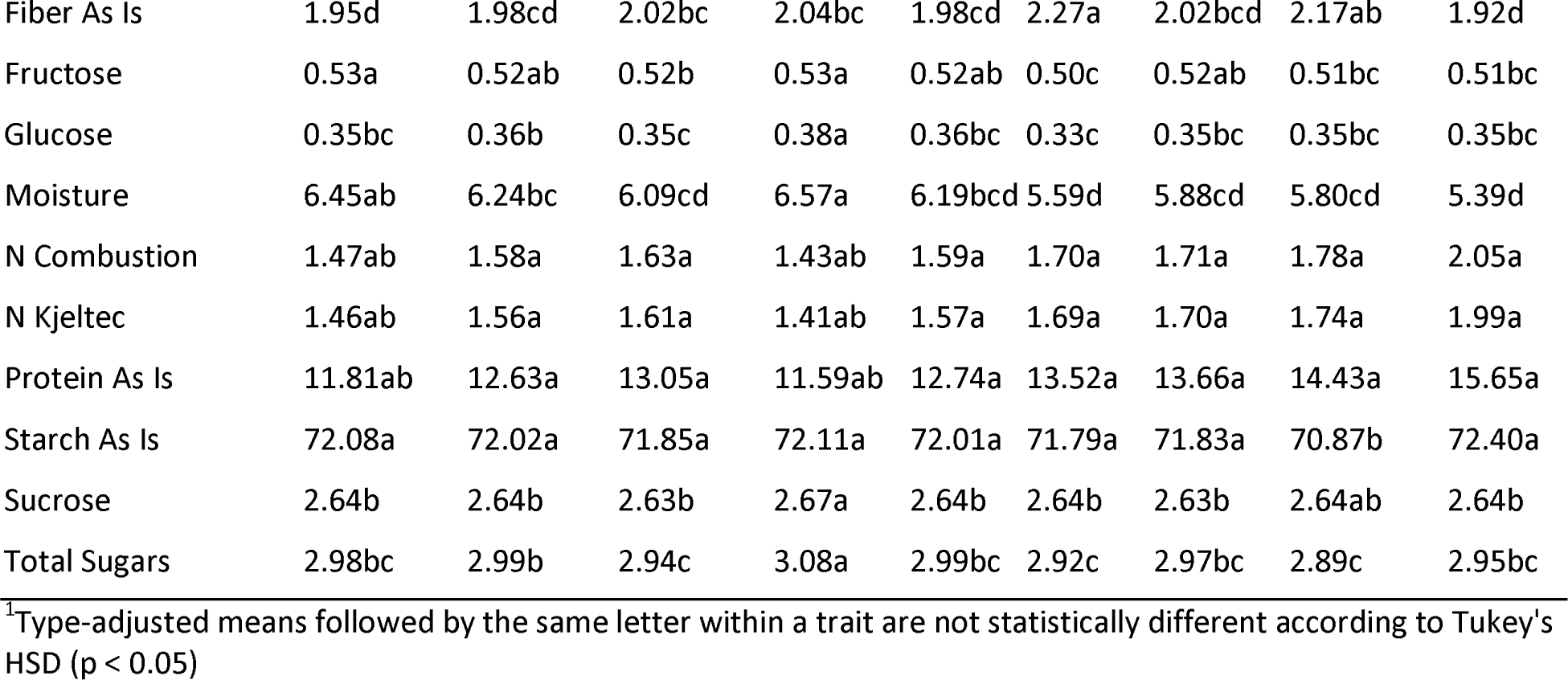
Summary of best linear unbiased prediction (BLUP)-adjusted mean differences for each compositional trait compared among maize types.

### Genetic and environmental factors account for most of the variation in compositional traits

Given the high degree of variation observed for all of the compositional traits, we next sought to determine the sources of the variation. The proportion of variance explained by each term from the random effects model was analyzed, and the environment term, which combines year and location, was statistically significant (p-value < 0.001) for all traits and accounted for 3.26 to 81.68% of the total observed variation, with a mean of 40.44% (Figure 2; Supplemental Table 5). The environment term explained the majority of the phenotypic variance for ash, crude fat, crude fiber, fat as is, fiber as is, fructose, glucose, moisture, starch as is, and total sugars. To investigate the role of a location or year effect on compositional traits, the environment term was partitioned into location and year effects within the random effects model. The location effect ranged from 0 to 61.60%, the year effect ranged from 0 to 32.18%, and the location by year interaction ranged from 0.01 to 85.33% (Supplemental Figure 6). The location effect was the predominant source of variation for ankom crude fiber, ash, crude fat, crude fiber, fat as is, fiber as is, fructose, and starch as is (Supplemental Figure 6). The year effect explained the most phenotypic variance for glucose and total sugars. The interaction between location and year was the largest for moisture, nitrogen combustion, nitrogen Kjeltec, and protein as is. For many of the carbohydrate compounds analyzed the location was the driving factor of the environmental effect, which can also be seen in the distribution of the raw phenotypic data (Figure 1). Of the five unique environments, the MO 2017 environment, which is the most geographically and climatically distinct environment, was the lowest for all carbohydrate traits. Previous studies have also found a statistically significant effect of the environment on compositional traits. For instance, oil concentration in the grain has been shown to be greatly influenced by geographic location and planting year (Dunlap et al., 1995; Jellum & Marion, 1966). Likewise, a study of seven maize hybrids grown in Europe analyzing compositional traits, found 1,986 statistically significant differences out of 4,935 comparisons between the four distinct environments included in the study, highlighting the importance of environment effects for these traits (Reynolds et al., 2005). These results highlight that environmental factors, such as location and year effects, need to be taken into consideration for breeders to meet breeding targets for compositional traits.

**Figure 2.**
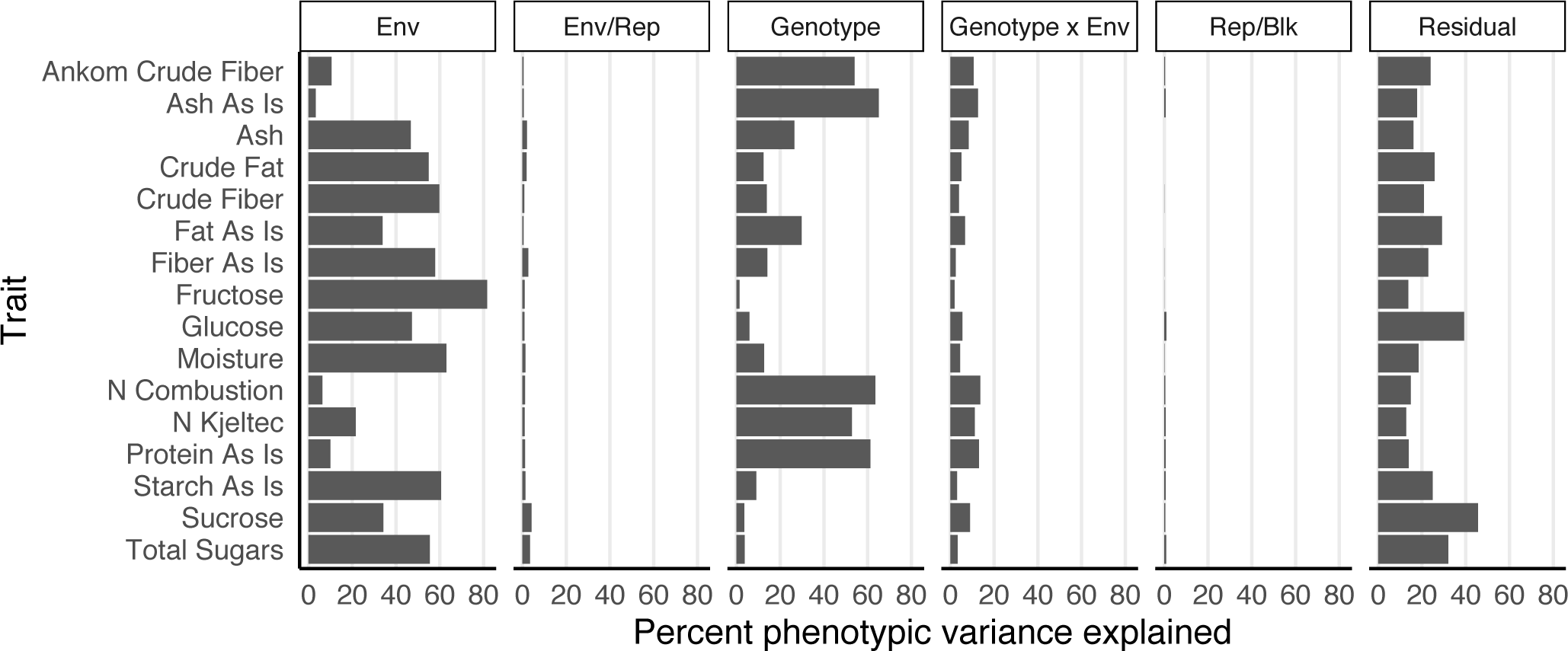
Percent of phenotypic variance explained by each factor from the ANOVA of the random effects model for sixteen compositional traits collected from the WiDiv association panel. Env, environment, Rep, replication, Blk, block.

The genotype and genotype by environment interaction terms were also statistically significant for all traits (p < 0.001) and ranged from 1.43 to 65.09% (average 26.87%) and 1.95 to 13.68% (average 7.18%), respectively (Figure 2; Supplemental Table 5). In contrast, a previous study analyzing a different maize association panel found kernel moisture and starch content had a non-significant genotype by environment interaction, while oil and protein content did have a statistically significant genotype by environment interaction (Flint-Garcia et al., 2005). However, in that study the authors noted that many environments had only a single replication, which caused the genotype by environment interaction to be confounded with the residual term in the mixed model which would affect significance testing. While the genotype by environment interaction was not the main source of phenotypic variation, it was still a statistically significant source of variation for all traits. Breeders have options for dealing with genotype by environment interactions: choose to ignore these interactions and instead choose varieties that are superior when averaged across environments, try to reduce these effects by testing varieties in similar environments, or try to exploit these interactions by selecting varieties that excel in a particular environment (Bernardo, 2010). For Ankom crude fiber, ash as is, nitrogen combustion, nitrogen Kjeltec, and protein as is, the genotype term in the model explained the most phenotypic variation, with 52.78 to 65.09% of the variation explained. While for ash, crude fat, crude fiber, fat as is, fiber as is, and moisture intermediate values of variation explained by genotype were observed ranging from 12.35 to 29.83% or low values for the carbohydrate traits ranging from 1.43 to 9.09% (Figure 2). The approach for dealing with genotype by environment interactions will likely need to be different for these different classes of traits.

Finally, the residual error ranged from 12.08 to 45.63% of variance explained and accounted for the highest variance explained for sucrose, which had only 34.21% of the variation explained by genotype (Figure 2). The high residual variance for some traits may reflect the lower quality of the local prediction equations for those traits (Supplemental Figure 3), and also the detection limits of analytical techniques specifically in the case of total sugars, glucose, fructose, and sucrose (<0.15%; Supplemental Table 2). Still, a sizable proportion of variance explained was due to residual variation for the rest of the compositional traits. Some of these factors could be epistatic effects or micro-environmental factors that were not taken into consideration in our random effects model.

### Heritabilities of compositional traits and implications for single trait breeding targets

Trait heritability is an important consideration in determining the ability to make gains from selection for a particular trait by a breeder. Traits that have a high heritability have better response to selection (Falconer & Mackay, 1996), and understanding the heritability for compositional traits in the breeding population helps breeders plan where to invest or divest resources to make genetic gains for the desired breeding target. Broad sense heritabilities based on entry-means for the diversity panel ranged from 0.36 (sucrose) to 0.94 (ash as is; protein as is; nitrogen combustion; nitrogen Kjeltec) (Table 1). With the exceptions of fructose, glucose, sucrose, and total sugars, the compositional traits measured in this study all had relatively high heritabilities (>0.75) (Table 1), indicating a larger genotypic variance compared to genotype by environment and residual variances that were measured in multiple environments. A previous study evaluating heritability in a panel of diverse inbred lines (Flint-Garcia et al., 2005) also found high heritabilities for protein, oil, and starch from NIR predictions that ranged from 0.83 to 0.91 (Cook et al., 2012). The lower heritabilities seen in the sugar-related traits in our study are likely a reflection of a lower quality prediction model (r = 0.260 to 0.649; Supplemental Figure 1) as compared to the global predictions equations.

The observed variation for compositional traits within the WiDiv association panel (Figure 1) and the relatively high heritabilities (Table 1) are critical for breeders to make gains from selection for quality traits within their breeding programs. The high heritabilities for these NIR predictions indicate that NIR is a suitable, high throughput, and affordable method compared to wet lab techniques to quickly evaluate germplasm within a breeding program and obtain robust data for making breeding decisions. Ten of the sixteen traits showed that the environment was the predominant source of phenotypic variation in the random effects model. For these traits, potential improvement could be made by breeding for target environments or growing and sourcing from certain environments that are more favorable for specific traits of interest. Ankom crude fiber, ash as is, nitrogen combustion, nitrogen Kjeltec, and protein as is had the majority of the phenotypic variation partitioned to the genotype term in the random effects model, and had very high heritabilities with an average of 0.94 (Table 1). For these traits, breeders can make genetic gains through selection that can be deployed across more diverse environments.

### Correlations between compositional traits among types of maize and implications for multitrait breeding

Statistically significant phenotypic correlations were detected among the WiDiv association panel as a whole with only nine non-significant correlations (Supplemental Table 6). The strongest correlations were between the various protein and nitrogen measurements, including nitrogen Kjeltec, nitrogen combustion, and protein as is (r = 0.99-1.00). Nitrogen combustion is collected via the Dumas method (Dumas, 1831), which determines total nitrogen content (organic and inorganic nitrogen) by combusting a sample at a high temperature in an oxygen-rich environment to convert the nitrogen in the sample into nitrogen gas. Nitrogen Kjeltec is measured with the Kjeldahl method (Kjeldahl, 1883) that determines the total organic nitrogen content of a sample through a sulfuric acid digestion. Protein as is is then calculated by a protein conversion factor of 6.25 for maize to estimate the total protein content in a sample. The fact that these traits are highly correlated is not particularly surprising as they are measuring the same compound in different ways, but gives confidence to the quality of the data.

By investigating phenotypic correlations of distinct compounds, we found that protein as is was negatively correlated with starch as is (r = -0.23, p-value < 0.001) (Supplemental Table 6). The negative trade-off between protein and starch throughout the history of maize breeding is well known. Breeding for increased yield in maize has driven an increase in starch content over time with an overall decrease in protein content (Duvick & Cassman, 1999). This is because a lower energy threshold is required of the plant to make starch since it is a primary product of photosynthesis, while protein requires more energy and relies on external factors such as nitrogen availability in the soil (Jenner et al., 1991). In a previous study by Cook et al., (2012) a negative correlation was found between NIR-predicted starch and protein (r = -0.56) in a panel of 282 diverse inbred lines. We also found a negative phenotypic correlation between starch and fat (r = -0.47) and a positive correlation between protein and fat (r = 0.52), both of which were consistent with Cook et al., (2012). In the maize kernel, fat content is primarily found in the germ, and starch and protein are found in the endosperm. The endosperm is roughly divided into the floury endosperm that contains mostly starch and which is located in the interior of the kernel, and the vitreous endosperm surrounding it and which contains much of the protein found in the kernel (Holmes et al., 2019). For the purpose of food-grade maize, a shift in the proportion of floury endosperm to vitreous endosperm, or a larger germ relative to the endosperm, will be needed to increase the relative amount of protein and fat in the kernel.

Breeding efforts for different applications (e.g. food-grade corn, sweet corn, popcorn, etc.) are focused on different needs based on the product market, which have resulted in very different trait means (Table 2), and potentially different correlations among traits within these groups of germplasm. Pearson correlations were recalculated after separating the lines into dent (Stiff Stalk, Non-Stiff Stalk, and Iodent) versus popcorn, sweet corn, and flint since these two groups showed the most statistically significant differences in BLUP-adjusted means between traits (Table 2). Overall, more statistically significant pairwise correlations were detected for the dent germplasm compared to popcorn, sweet corn, and flint (Figure 3), likely in part due to the substantially larger sample size for the dent group. The overall pattern of pairwise correlations were relatively similar, but the changes in magnitude and/or direction were substantial in some instances (Figure 3). Starch as is had weak positive correlations with glucose, fructose and sucrose (r = 0.36, 0.23, and 0.13) in pooled dent germplasm, but in popcorn, sweet corn, and flint lines these correlations changed direction and were stronger negative correlations (r = -0.65, -0.61, and -0.57) (Figure 3). Similar changes in correlation patterns were also observed after separating lines into dent and non-dent for correlations between crude fat and numerous other traits (ankom crude fiber, ash, fructose, nitrogen Kjeltec, and nitrogen combustion), as well as the correlation of crude fiber with both moisture and protein. The correlation between protein and crude fat was positive (r = 0.28) in the dent germplasm, but was reversed to a negative relationship (r = -0.67) in the non-dent germplasm.

**Figure 3.**
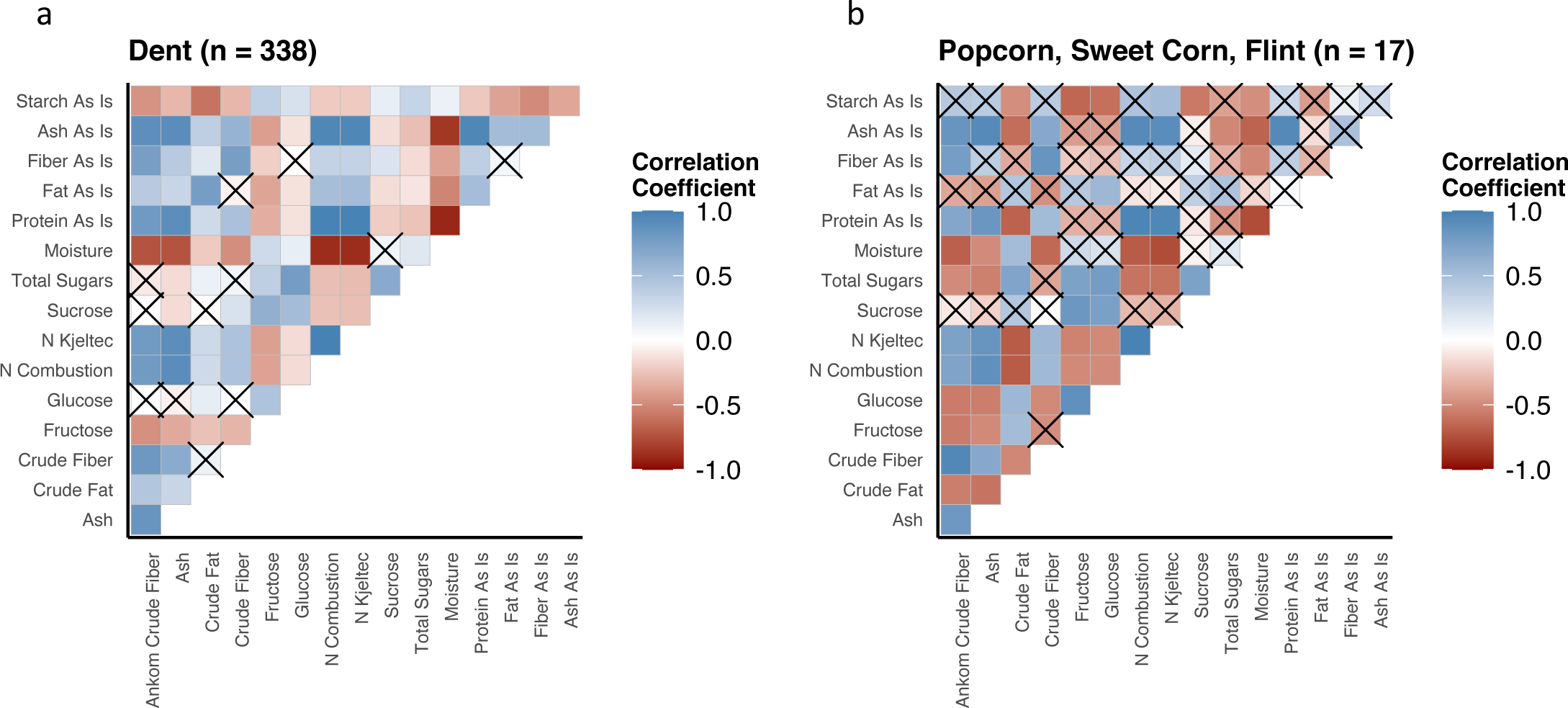
Heatmaps of Pearson correlation matrices for compositional traits partitioned by a) dent, b) popcorn, sweet corn, and flint. Squares that are crossed out represent non-significant Pearson correlations (p-value > 0.05).

Breeding for a single compositional trait can be a difficult task, given controlling genetic and environmental factors, but when improving multiple traits simultaneously in a breeding program the task becomes even more complex and difficult. Phenotypic correlations in the full WiDiv association panel and parsing the panel into dent and non-dent types revealed that there are many phenotypic trade-offs that need to be assessed by a breeder in order to make desired genetic gains. For instance, if a breeder has a program goal of improving quality, certain phenotypic correlations may work for or against the breeding target. Protein in the dent germplasm pool was negatively correlated with starch (r = -0.23), but positively correlated with ash, fiber, and fat (r = 0.96, 0.37, and 0.51) (Figure 3). Selections for increasing protein content would then also translate to gains in ash, fiber, and fat but a decrease in starch content that would meet the breeding targets of food-grade maize. These compositional targets help with physiological aspects of the kernel such as kernel damage, stress cracks, and higher protein content that impact processing. There are a few methods for multiple trait selection such as tandem selection, independent culling, and index selection (Bernardo, 2010). In tandem selection, a breeder selects for a single trait at a time until breeding targets have been met. This method can be time consuming, but can be efficient if traits are correlated with each other in a way that meets the breeding targets. Independent culling on the other hand selects for multiple traits in a single breeding cycle with minimum threshold levels determined for each trait, and entries that are above these thresholds continue through the breeding pipeline. Index selection selects for multiple traits simultaneously using an index value determined by the breeder. The index can weight traits based on economic value alone, genetic and phenotypic covariances with economic values, and performance (Elston, 1963; Hazel, 1943; Smith, 1936; Williams, 1962). Index selection is a more effective method than both tandem selection and independent culling while having the ability to have multiple traits imputed into an index. Index selection can also be tailored to a breeding program with multiple methods from which a breeder can choose. For instance, a food-grade maize breeder with the goal of increased quality would put higher weights on protein, fiber, and fat content with a smaller weight on starch content. Knowledge of how compositional traits interact with each other at a phenotypic level is important for making genetic gains in a breeding program, and understanding the genetic architecture of these traits can help accelerate breeding goals for improved quality in maize grain.

### Compositional traits are mostly controlled by a large number of small effect loci

To determine the genetic architecture of our compositional traits we utilized a GWAS approach, as this population was originally developed for this purpose (Hansey et al., 2011). For this analysis we utilized a set of 2,386,666 SNPs that had been previously called from whole genome resequencing data (O’Connor et al., 2020). Principal component analysis (PCA) using these SNPs showed clear separation of the three major maize dent heterotic groups: Stiff Stalk, Iodent, and Non-Stiff Stalk (Figure 4a). The pattern observed in the PCAs are similar to other published results (Romay et al., 2013; van Heerwaarden et al., 2012; White et al., 2020) describing the separation of maize types. The clear separation of maize types from the WGS data gave us confidence in the genotypic data, and provided covariates to use in the GWAS to account for population structure.

**Figure 4.**
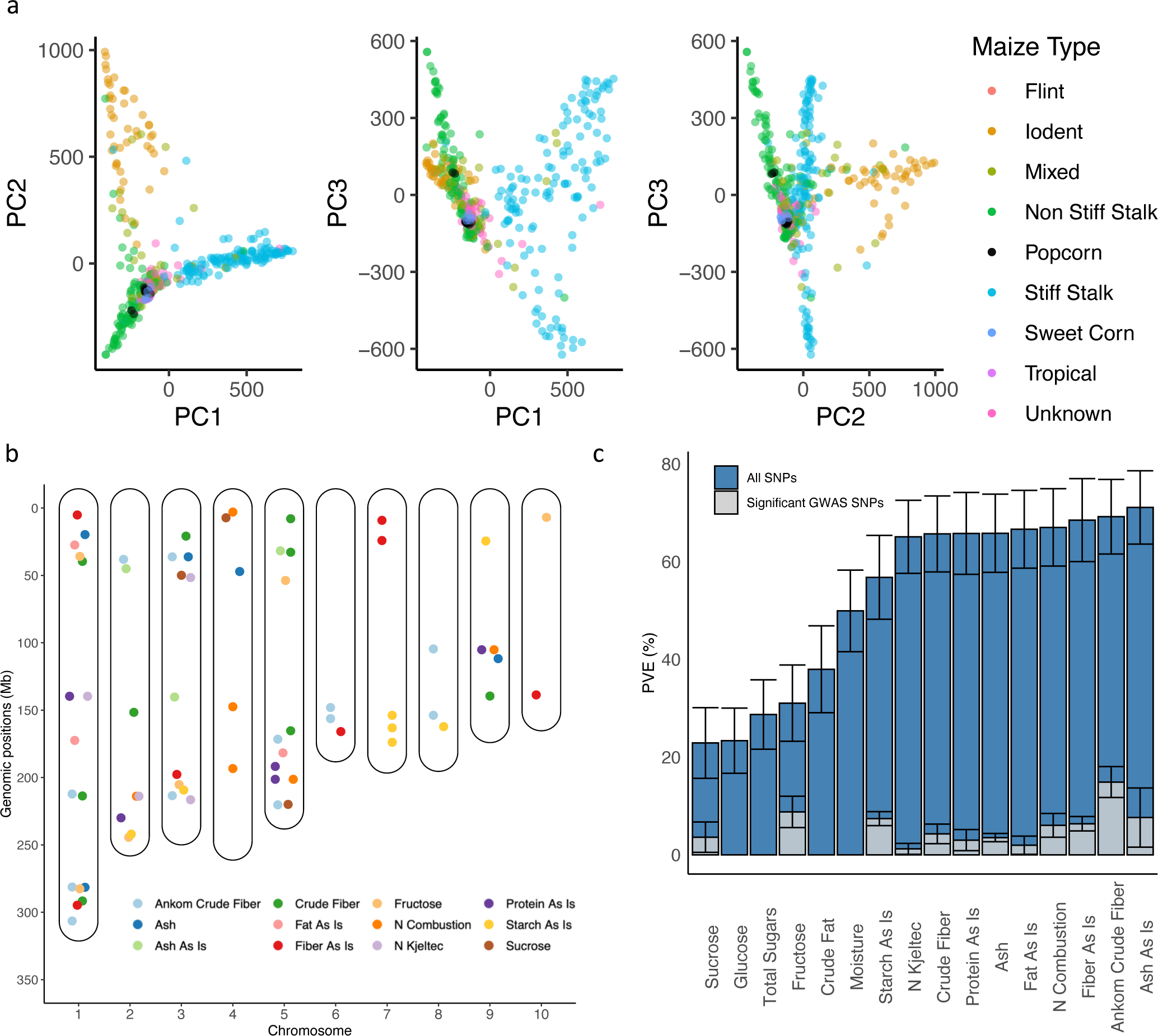
Genetics architecture of compositional traits in a maize diversity panel. a) Pairwise plots of the principal components of population structure using 2,386,666 SNPs for the 446 inbred lines used for genome-wide association studies (GWAS). Each point is colored based on maize type classification. b) Statistically significant GWAS SNP associations of compositional traits. Each point represents the physical location of a statistically significant SNP on the chromosome. Points are colored corresponding to a particular compositional trait. c) Percent variation explained (PVE) by the full set of 2,386,666 SNPs used in the GWAS are colored in blue, while the gray corresponds to significant SNPs that were specific for each trait based on GWAS results. Error bars represent the standard error.

For all 16 compositional traits in this study, the genetic architecture was mapped using 446 inbreds with resequencing data using the MLMM in FarmCPU (Liu et al., 2016) with 2,386,666 SNPs. A total of 70 statistically significant loci across 12 of the 16 traits were detected based on a genome-wide corrected Bonferroni threshold of -log10(p) = 7.68 (Figure 4b, Supplemental Figure 7, Supplemental Table 7). The traits with statistically significant GWAS loci included Ankom crude fiber (12), ash (5), ash as is (3), crude fiber (9), fat as is (3), fiber as is (7), fructose (6), nitrogen combustion (6), nitrogen Kjeltec (4), protein as is (5), starch as is (7), and sucrose (3) (Figure 4b). In a previous study conducted by Cook et al., (2012) using NIR predicted data for starch, protein, and oil, no significant associations were found using a MLM method for GWAS, which is contrary to our study in which starch, protein, and fat from the global equations had statistically significant loci. The differences between the two studies could be due to a few factors such as having more individuals in the WiDiv association panel, a more balanced allele frequency profile because the WiDiv is less genetically diverse compared to the 282 panel, a much larger SNP dataset (43 times as many markers), and a more powerful GWAS model to detect significance for these complex compositional traits.

As we had multiple traits in this study that measured the macro-molecules in different ways for protein, starch, and ash, we decided to first look at the overlap in hits within these sets of traits. There were four instances of nitrogen-related compositional traits sharing the same associated SNP (Figure 4b, Supplemental Table 7): chromosome 1 between nitrogen Kjletec and protein as is, chromosome 2 between nitrogen combustion and nitrogen Kjeltec, and chromosomes 5 and 9 between nitrogen combustion and protein as is (Figure 4b). The shared statistically significant loci between nitrogen combustion, nitrogen Kjeltec, and protein as is are not unexpected as these traits are highly correlated (r = 1.00, r = 0.99, r = 0.99) with each other and are measuring nitrogen content in slightly different fashions, as described above (Supplemental Table 6). While the nitrogen-related compositional traits shared many SNP hits, the fiber and carbohydrate related traits all gave different statistically significant loci. This is not particularly surprising as fiber can be measured in multiple ways, reflecting different compounds within the fiber fraction (Mertens, 1992). This is exacerbated when comparing the local equations to the Perten global equations. The local equations were developed with a single method, while the global equations were developed using a mixture of methods that integrated data from different laboratories (D.E. Honigs, Perten Instruments, personal communication), which may reduce signal gained from using equations developed from a single method and measuring a specific aspect of fiber.

Across the 70 statistically significant SNPs, 47 were located within a gene in the B73v4 annotation, and the remaining 23 were within 2 kb of a gene model (Supplemental Table 7). Functional annotations for the genes nearest each statistically significant SNP were determined by using a series of databases (see Materials and Methods). Only 11 statistically significant loci yielded uncharacterized protein annotations from the combined gene annotation methods (Supplemental Table 7). Some notable associations for Ankom crude fiber include 1) a transcription factor, *Dof4*, on chromosome 3, which regulates shoot branching and seed coat formation in *Arabidopsis thaliana* (Zou et al., 2013); 2) *Apk1* on chromosome 6, which encodes a protein kinase found to phosphorylate many proteins involved in the glycolysis pathway (Hirayama & Oka, 1992), and 3) a WUSCHEL-related homeobox gene on chromosome 8, which has previously been shown to be associated with maize embryo development (Nardmann et al., 2007). For fat as is, a bzip transcription factor 111 previously found to be involved in the carbon/nitrogen balance regulation (Wu et al., 2019) was identified as a candidate gene. While these results provide an exciting resource to breeders to make quality gains in maize, and this study identified more statistically significant SNPs than previous GWAS studies of compositional traits in maize (Cook et al., 2012), the loci identified in this study do not explain the full breadth of phenotypic variation in this population.

To gain more insight into the underlying genetic architecture of these compositional traits, we estimated the proportion of variance explained (PVE) by the full set of 2,386,666 SNPs generated from the WGS. The percent of PVE ranged from 22.90% to 71.09% with sucrose having the smallest amount and ash as is having the most (Figure 4c). PVE was also calculated for just the significant SNPs identified for each trait using only those SNPs that were significant in the GWAS for the trait. The PVE for these significant GWAS SNPs was substantially lower and ranged from 1.23% to 14.90% with nitrogen Kjeltec having the smallest PVE from the significant SNPs and Ankom crude fiber having the most (Figure 4c). The carbohydrate traits tended to have the lowest amount of variation explained ranging from 22.90% to 56.79% (Figure 4c) compared to the other traits. This could be due to the accuracy of the NIR prediction equations used to generate values for the sugars or reflects the large role that environment contributes to these traits (Figure 2). Nine of the 16 traits had relatively large (>60%) PVE attributed to the full set of SNPs, indicating that many loci of small effect contribute to variation of compositional traits and did not meet the threshold of significance rather than a few large effect loci identified in the GWAS. This result is consistent with results found in the Illinois Long-Term selection experiment in a QTL study for oil concentration, where it was estimated that more than 50 loci may be involved in controlling the trait (Laurie et al., 2004). The upper limits for protein and oil have not been reached in the Illinois Long-Term selection experiment, even after over 100 generations of selection (Dudley & Lambert, 2004), giving reason to believe many of these small effect loci can be used to make tremendous gains for improved quality grain for these and other compositional traits. This result brings more complexity to the challenge for breeding to improve for quality traits and the cost of marker data across many small effect loci. Rather, a recurrent selection program may be needed to reach a breeding target when considering resource allocation in limited budgets.

## CONCLUSIONS

In summary, we collected NIR compositional traits on a set of 501 inbred lines adapted to the Upper Midwest grown in five unique environments. The WiDiv association panel showed substantial variation for compositional traits and it was shown that genetic and environmental factors account for most of the phenotypic variation observed. Statistically significant differences in means among types of maize and correlations between compositional traits were found to reflect the diversity of maize types and the underlying anatomical and structural make-up of the kernels. A total of 2,386,666 SNPs derived from WGS data was used to conduct a GWAS on 446 individuals from the WiDiv association panel, which resulted in 70 statistically significant loci detected. The proportion of variance explained by all of the SNPs showed that many compositional traits are controlled by small effect loci. Insights into the underlying genetic architecture of these traits and how they are correlated will help breeders in implementing breeding strategies for quality enhancement breeding targets that have been largely neglected to date.

## SUPPLEMENTAL MATERIAL

**Figure S1**. Genetic distance comparison between shared inbreds for RNA-seq (Mazaheri et al., 2019) and WGS (O’Connor et al., 2020) data.

**Figure S2**. Dendrogram of the 446 inbred lines of the WiDiv association panel with 2,412,791 SNPs from WGS data.

**Figure S3**. Comparison of observed and predicted values for local near-infrared (NIR) spectroscopy prediction equations derived from a subset of 100 inbred lines with diverse spectra profiles.

**Figure S4**. Quantile-quantile (Q-Q) plots of the sixteen compositional traits for the random effects model. The red line represents the 1:1 y=x line.

**Figure S5**. Scree plot of eigen values for the principal components from the whole genome sequencing data used on the WiDiv association panel.

**Figure S6**. Percent of phenotypic variance explained by each factor from the ANOVA of the random effects model for sixteen compositional traits collected from the WiDiv association panel.

**Figure S7**. Manhattan plots from the GWAS of each compositional trait. Each point represents a single SNP and the solid red line represents the Bonferroni corrected p-value threshold to deem a statistically significant SNP at the 0.05 level (-log(p) = 7.68).

**Table S1**. Description of the inbred lines within the Wisconsin Diversity (WiDiv) association panel that were included in this study.

**Table S2**. Compositional wet lab trait observations used to develop local equations based on 100 spectrally diverse inbred lines from the WiDiv association panel and the predicted trait values from those equations.

**Table S3**. Phenotypic data for inbreds of the WiDiv association panel collected at five unique environments.

**Table S4**. Best linear unbiased predictions (BLUPs) for compositional traits of inbreds in the WiDiv association panel.

**Table S5**. Analysis of variance (ANOVA) from a linear model for 16 compositional traits collected from the WiDiv association panel.

**Table S6**. Pearson correlation coefficients between all traits (BLUPs) evaluated in the full WiDiv association panel.

**Table S7**. Statistically significant SNP locations and corresponding candidate genes identified through GWAS of 446 inbred lines from the WiDiv association panel for compositional traits.

## AUTHOR CONTRIBUTIONS

CNH, MYN, SFG, DE, AW, NA conceived this experiment. JSR, AMG, TJH conducted the experiments, JSR, CHO, PJM analyzed the data, JSR visualized the data, JSR, CNH wrote the original draft, all co-authors edited and approved the final manuscript.

## ACKNOWLEDGEMENTS

This work was funded in part by NSF IOS-1546272 to CNH and MDY-N, PepsiCo, Inc. to CNH, the Iowa Agriculture and Home Economics Research Station Project IOW03649 to MDY-N, and USDA-ARS base funds to SF-G. The authors acknowledge the Minnesota Supercomputing Institute (MSI) at the University of Minnesota for providing resources that contributed to the research results reported in this paper.

---

**Supplemental Figure 1.**
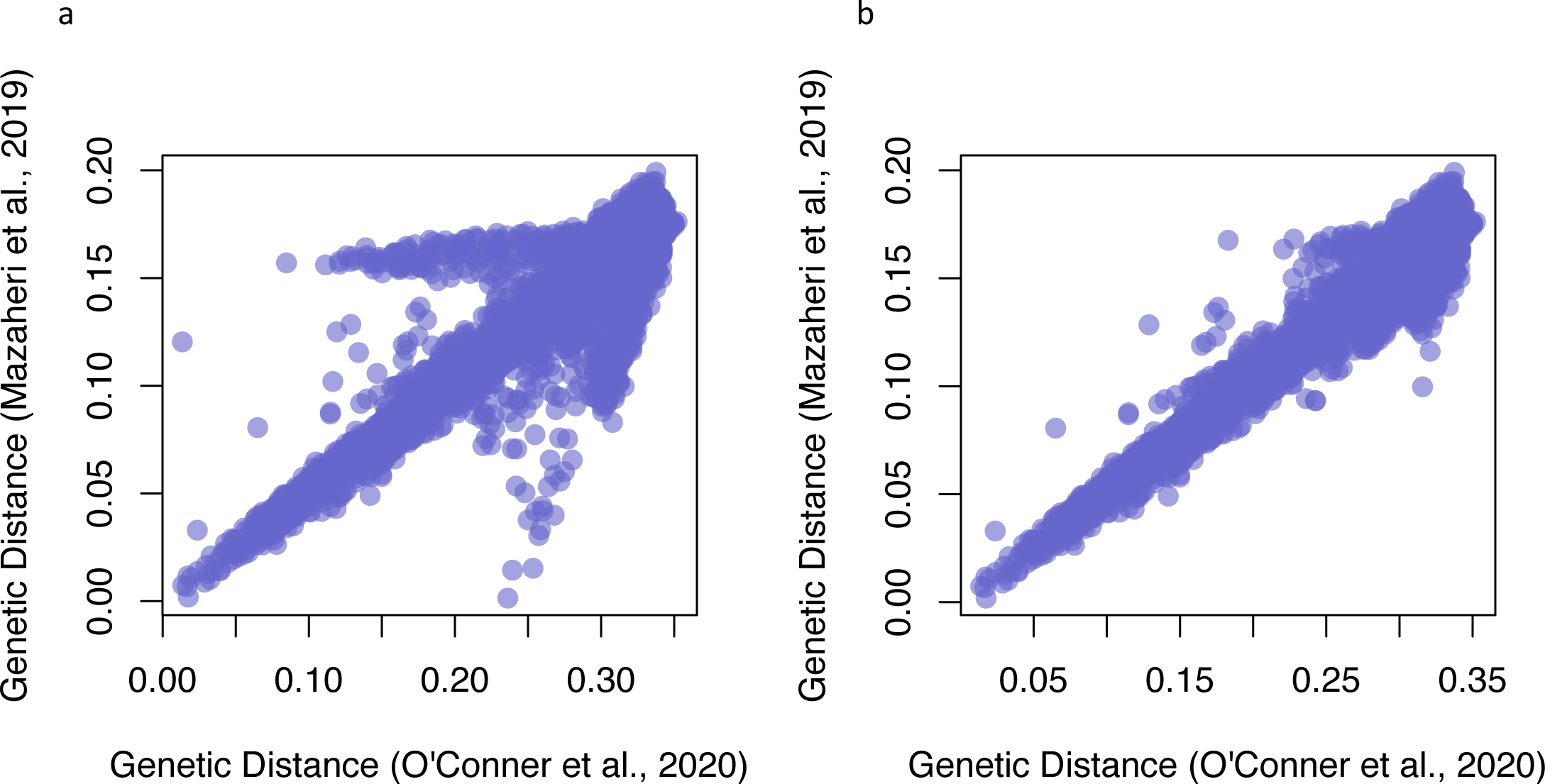
Genetic distance comparison between shared inbreds for RNA-seq (Mazaheri et al., 2019) and WGS (O’Connor et al., 2020) data. a) All inbreds included in GD comparison. b) Inbred lines B91, B14A, PHG86, and PHR55 removed due to inconsistencies between genotypic datasets.

**Supplemental Figure 2.**
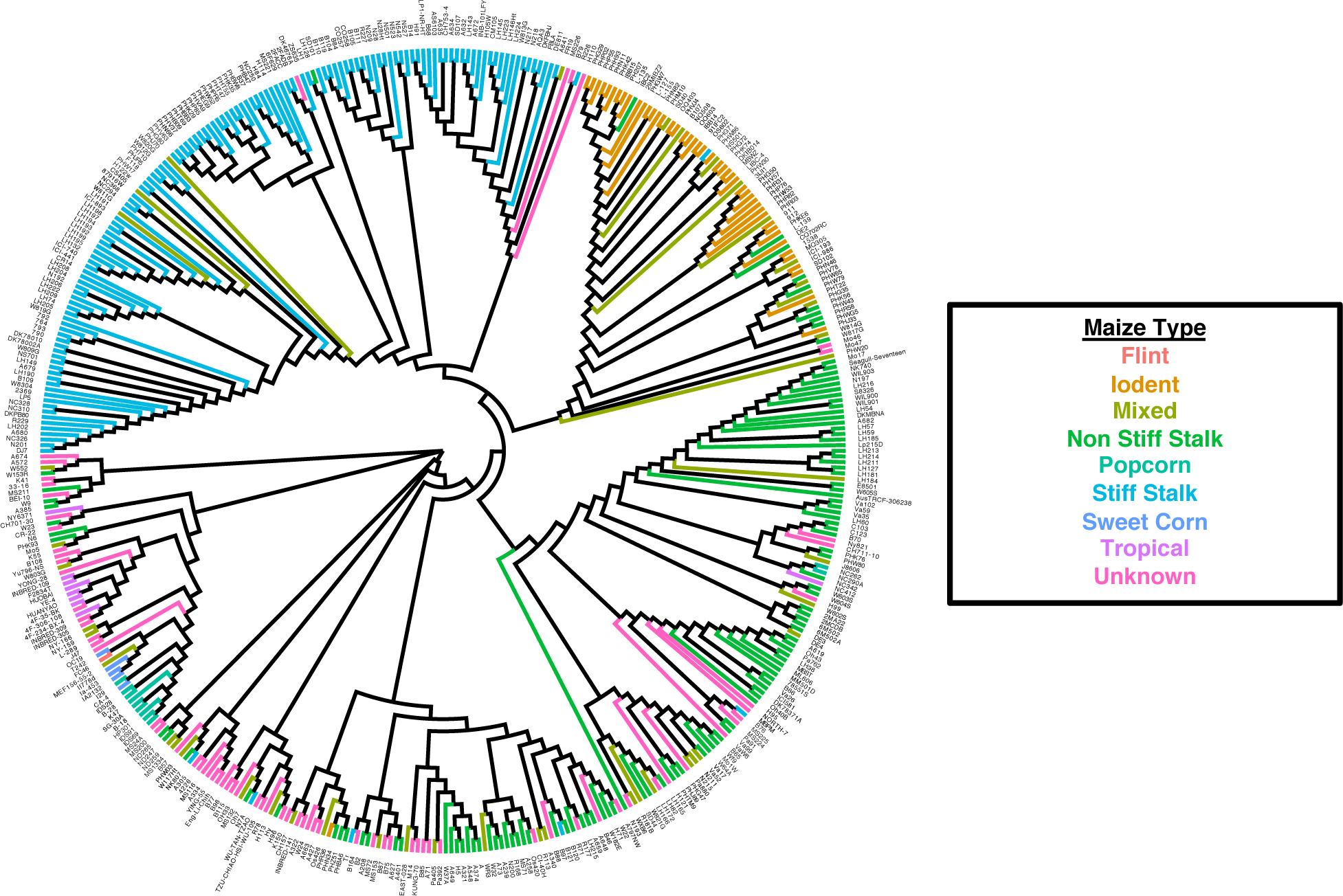
Dendrogram of the 446 inbred lines of the WiDiv association panel using 2,412,791 SNPs from WGS data. Distances were based on neighbor-joining clustering and inbreds are colored by maize type identification.

**Supplemental Figure 3.**
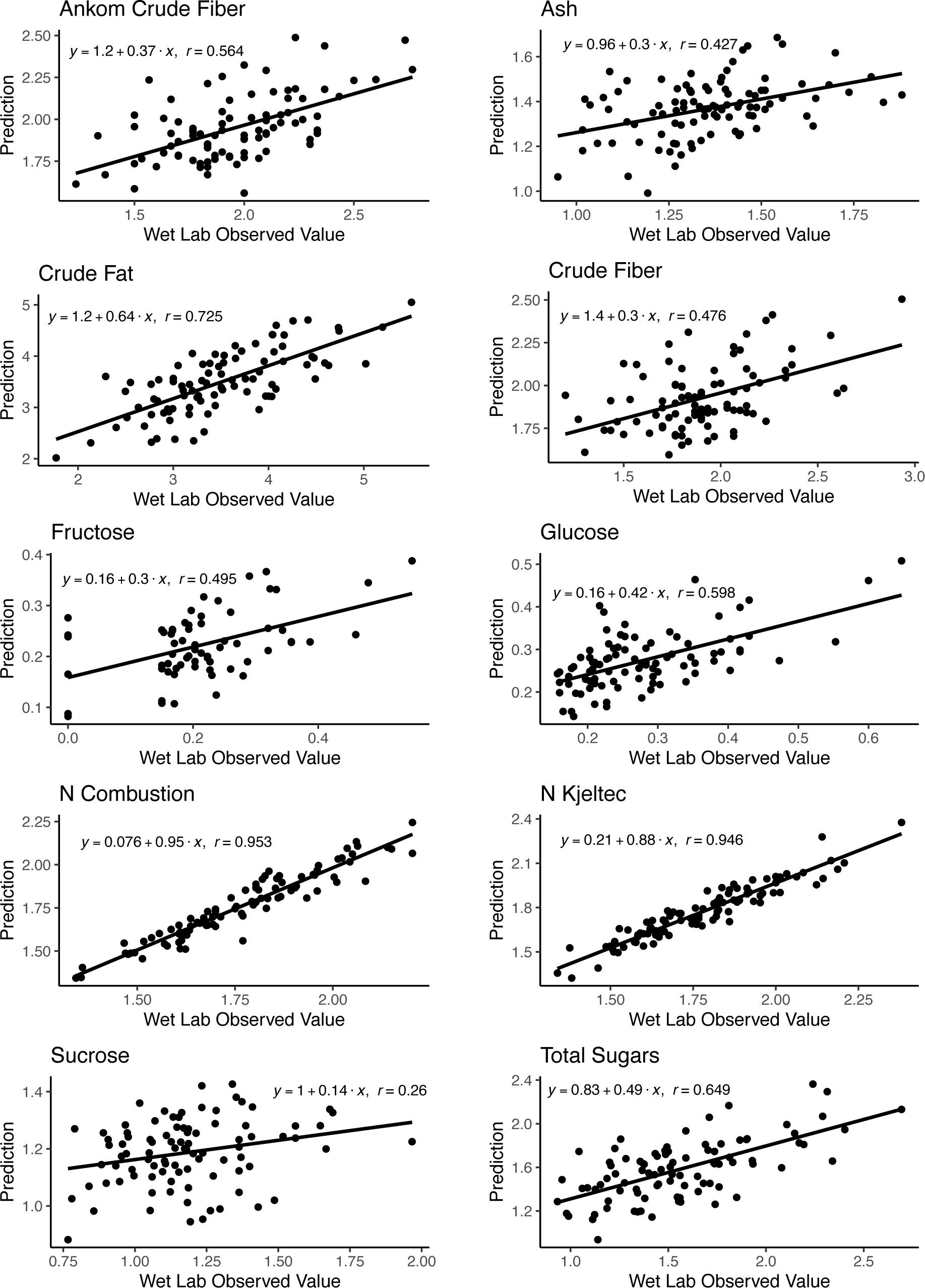
Comparison of observed and predicted values for local near-infrared (NIR) spectroscopy prediction equations derived from a subset of 100 inbred lines with diverse spectra profiles.

**Supplemental Figure 4.**
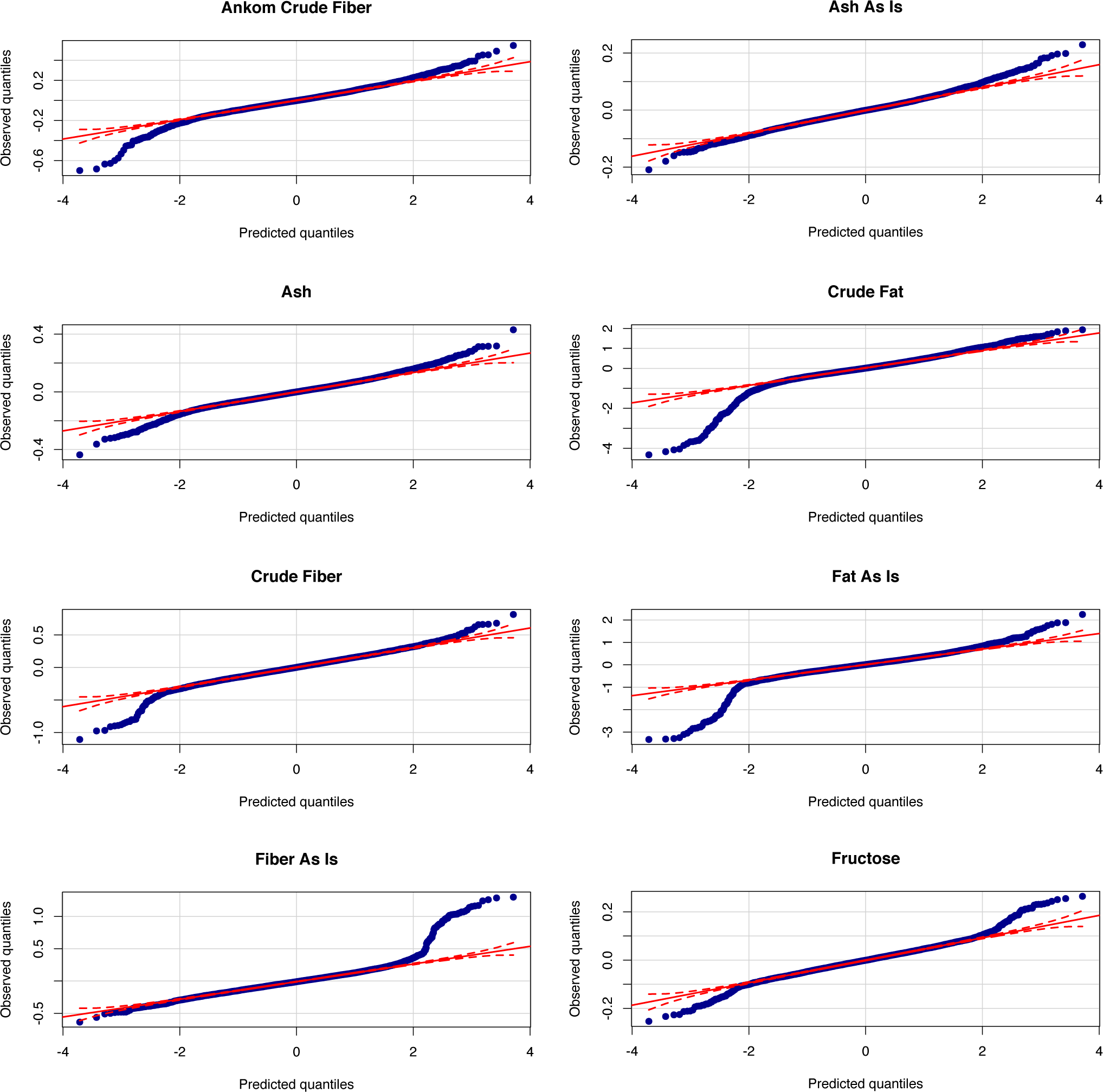
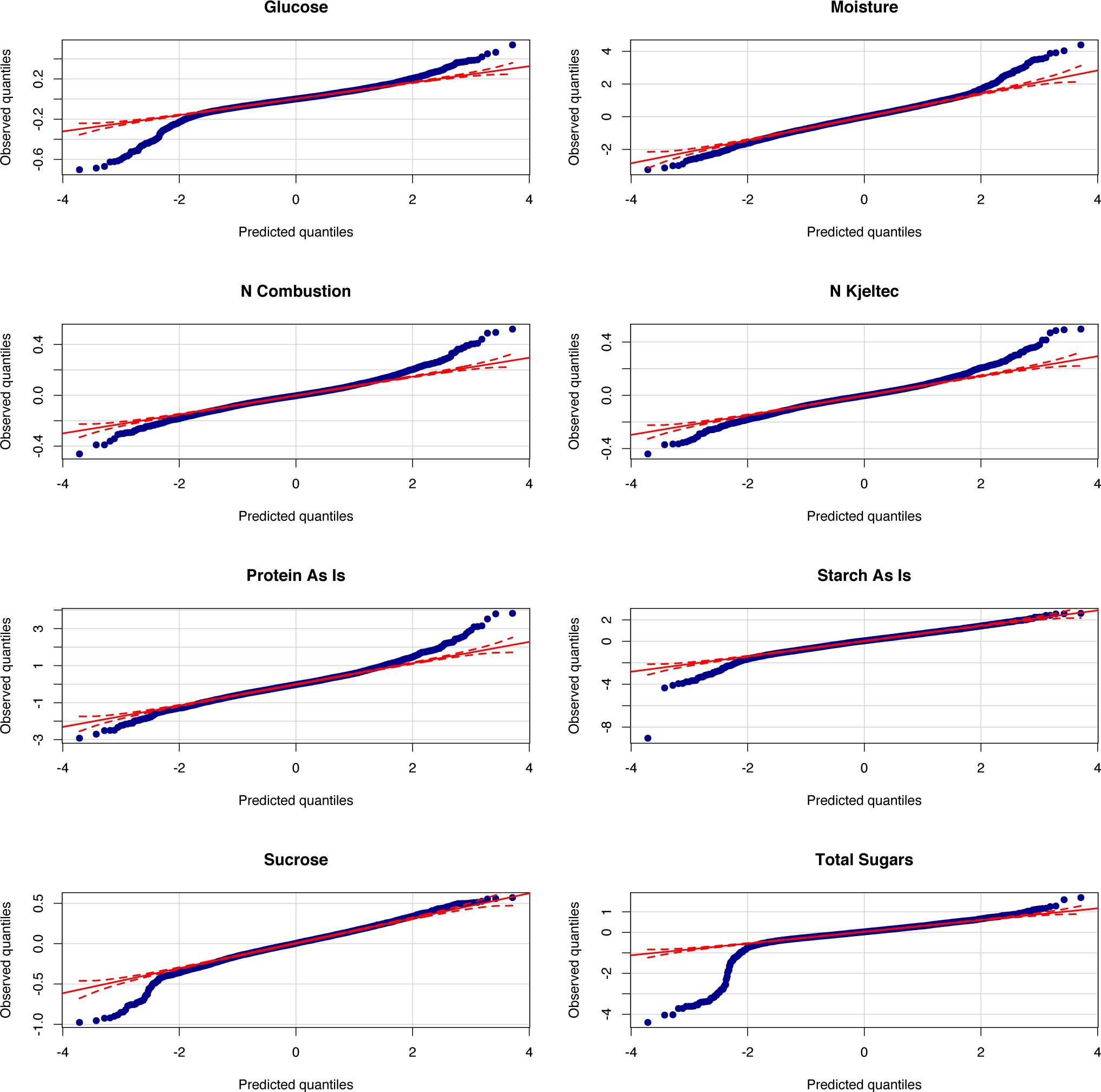
Quantile-quantile (Q-Q) plots of the sixteen compositional traits for the random effects model. The red line represents the 1:1 y=x line.

**Supplemental Figure 5.**
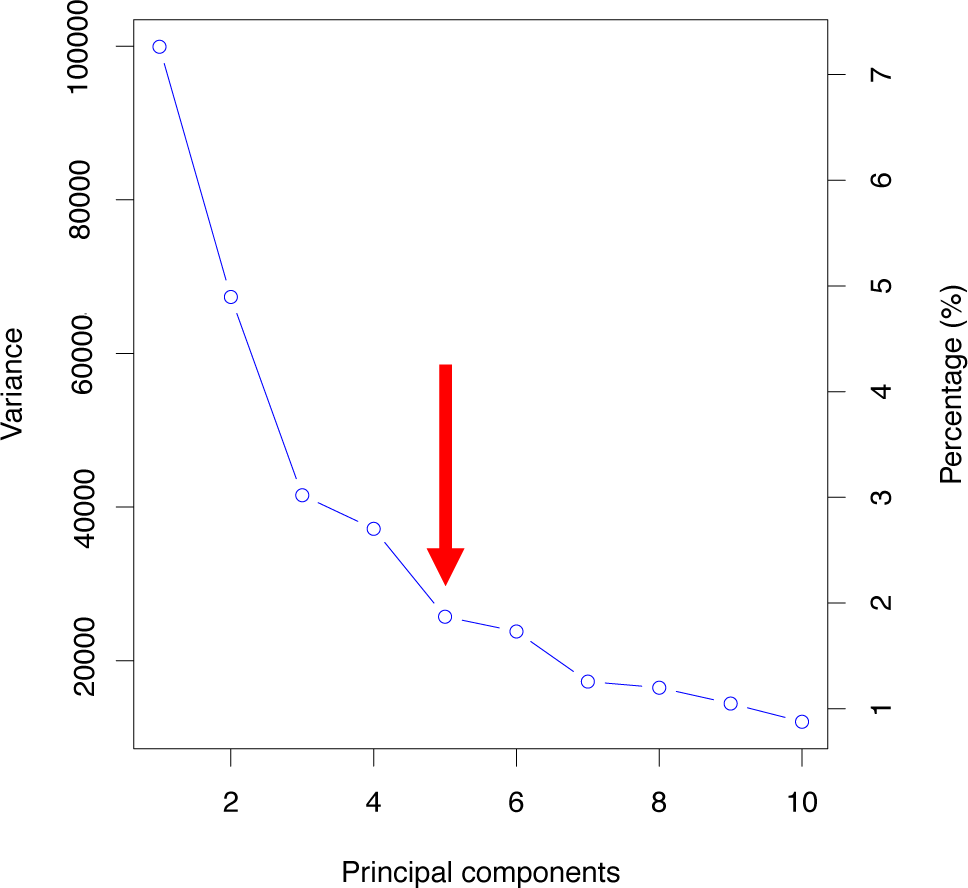
Scree plot of eigen values for the principal components from the whole genome sequencing data used on the WiDiv association panel. The red arrow represents the five principal components that were included as a cofactor in the FarmCPU model due to diminished variance explained for an additional principal components beyond n=5.

**Supplemental Figure 6.**
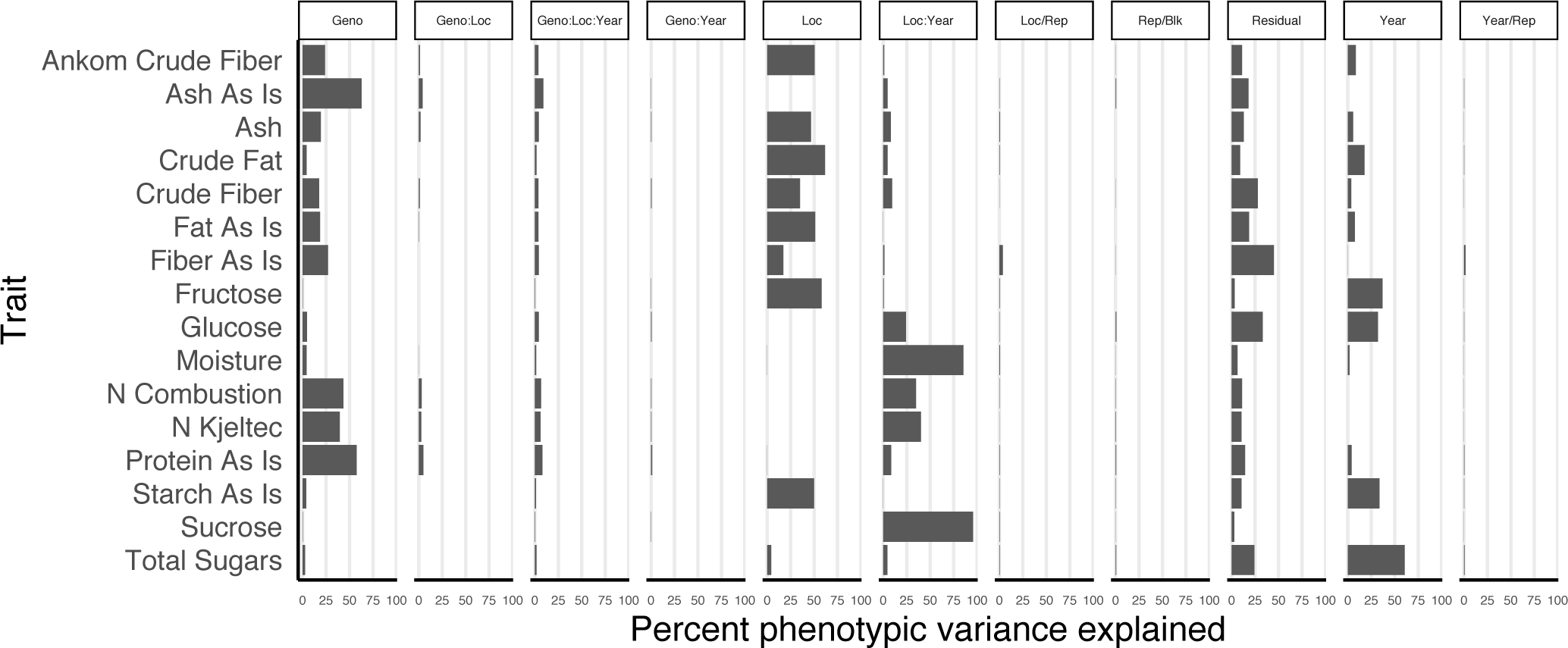
Percent of phenotypic variance explained by each factor from the ANOVA of the random effects model for sixteen compositional traits collected from the WiDiv association panel. Geno, genotype, Loc, location, Rep, replication, Blk, block.

**Supplemental Figure 7.**
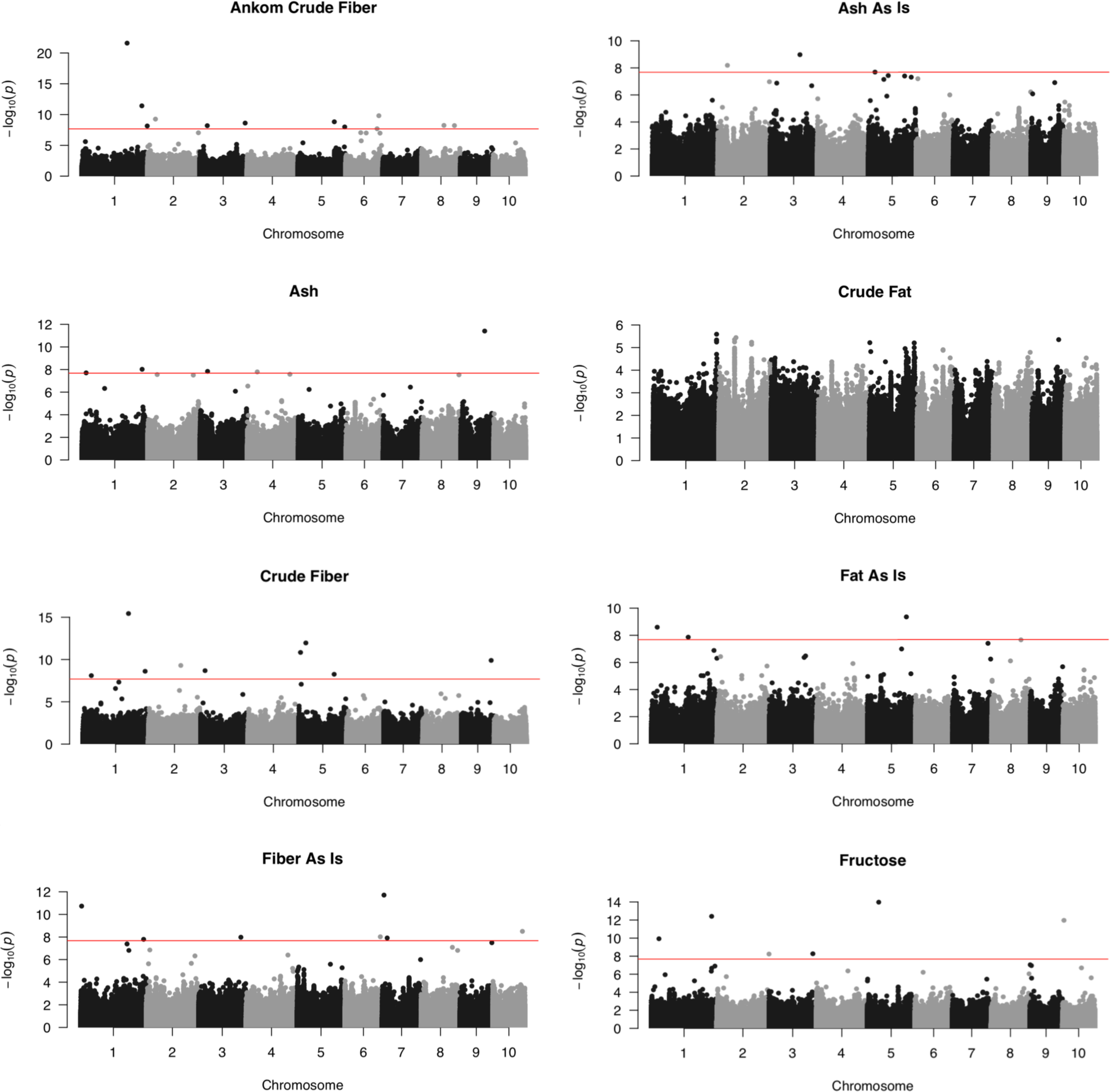
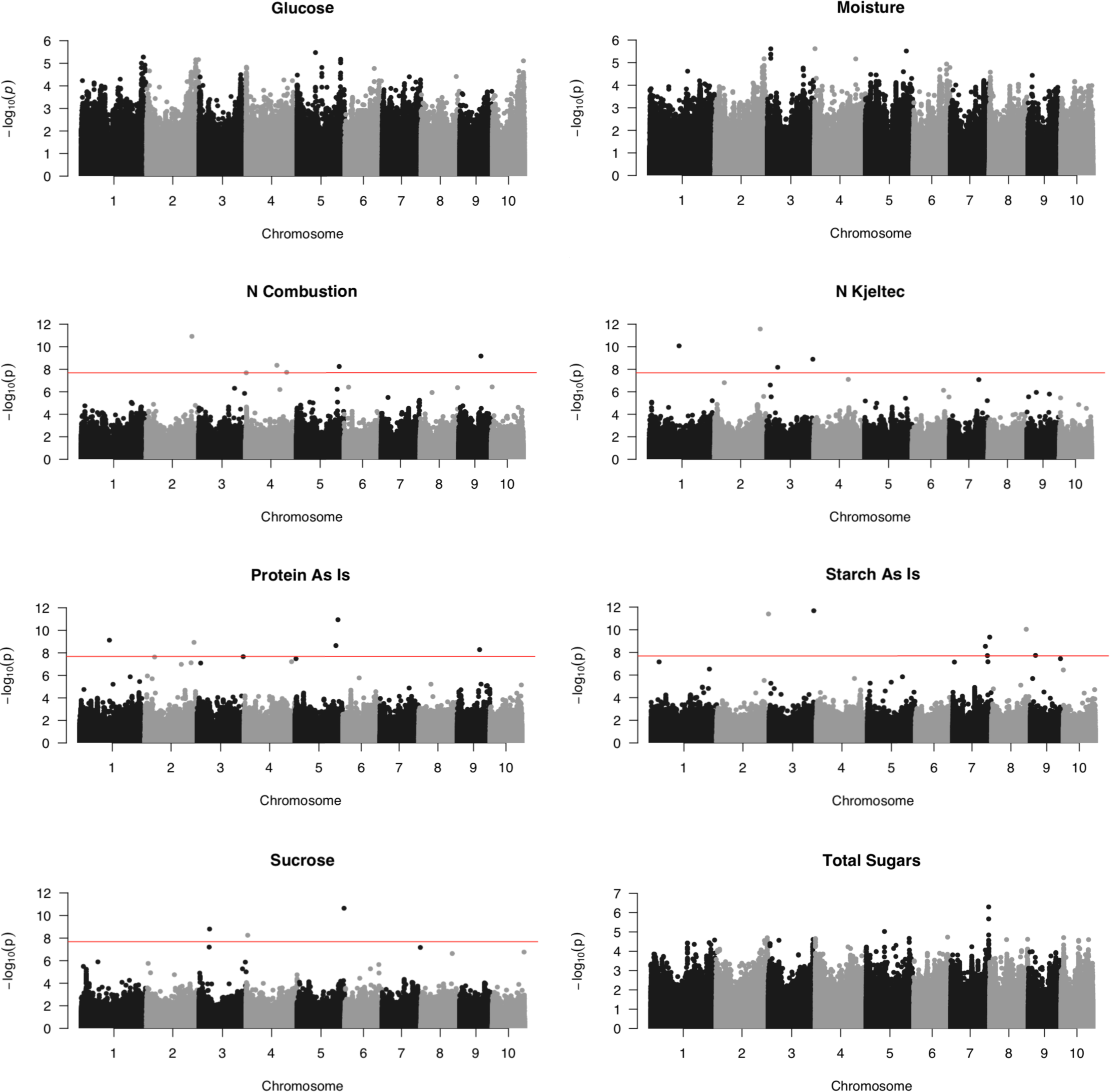
Manhattan plots from the GWAS of each compositional trait. Each point represents a single SNP and the solid red line represents the Bonferroni corrected p-value threshold to deem a significant SNP at the 0.05 level (-log(p) = 7.68).

